# An intrinsic cell cycle timer terminates limb bud outgrowth

**DOI:** 10.1101/296350

**Authors:** Joseph Pickering, Kavitha Chinnaiya, Constance A. Rich, Patricia Saiz-Lopez, Maria A. Ros, Matthew Towers

## Abstract

The longstanding view of how proliferative outgrowth terminates following the patterning phase of limb development involves the breakdown of reciprocal extrinsic signalling between the mesenchyme and the overlying epithelium (e-m signalling). However, by grafting mesenchyme cells from late stage chick wing buds to an early epithelial environment we show that this mechanism is not required. RNA sequencing reveals that mesenchyme cells terminate growth by an intrinsic cell cycle timer in the presence of e-m signalling. In this process, e-m signalling is required permissively to allow the intrinsic cell cycle timer to run its course. We provide evidence that a temporal switch from BMP antagonism to BMP signalling controls the intrinsic cell cycle timer during limb outgrowth. Our findings have general implications for other patterning systems in which extrinsic signals and intrinsic timers are integrated.

## Introduction

Detailed models of pattern formation have been derived in which cells are either informed of their position in the embryo by gradients of extrinsic signals, or by measuring intrinsic time. Recently, it has become evident that both processes often operate together - either in parallel, such as in antero-posterior patterning of the limb (thumb to little finger)^1^ ^2^ and in neural tube patterning^3^ - or sequentially, such as in proximo-distal patterning of the limb (shoulder to finger tips)^4–7^. Thus, understanding the precise requirement for extrinsic signals and intrinsic time in pattern formation is a challenging task. In addition, little is known about how these mechanisms operate together to terminate growth of tissues and organs.

During early stages of proximo-distal patterning of the chick wing (stage Hamburger Hamilton HH19), signals from the main body axis specify proximal identity (humerus)^4,8–10^. Once distal mesenchyme cells are displaced by growth away from proximal signals, a switch to an intrinsic timing mechanism occurs that specifies distal identity (elbow to digits)^6^. This intrinsic timer involves cells executing a programme of proliferation, expressing regulators of distal position (i.e. *Hoxa13*) and altering their cell surface properties, which are considered to encode position along the proximo-distal axis. In addition, distal limb bud mesenchyme cells maintain the overlying apical ectodermal ridge^11^ - a thickening of the distal epithelium, which is required for limb bud outgrowth until HH29/30^12,13^ (the final distal phalanx is laid down at around HH28). The apical ectodermal ridge maintenance factor is encoded by the product of the Bone Morphogenetic Protein (BMP) antagonist, *Gremlin1* (*Grem1*), which sustains expression of genes encoding Fibroblast Growth Factors (FGFs) in the apical ectodermal ridge^14–16^. The breakdown of the epithelial-mesenchyme (e-m signalling) feedback loop and loss of apical ectodermal ridge signalling is considered to terminate limb bud outgrowth at the end of the patterning phase (HH29)^17^ ^18^. This could suggest that a switch from an intrinsic timer back to extrinsic signalling occurs at later stages of proximo-distal patterning. Further growth of the limb is then a consequence of the expansion of osteogenic progenitor cells in the developing long bones.

However, we recently showed that grafts of old mesenchyme cells (HH27) expressing Green Fluorescent Protein (GFP) made to younger HH20 buds, which were left to develop for 24 h until HH24 (now referred to as HH24 graft-HH24g), intrinsically completed their proliferation programme in the presence of an overlying host apical ectodermal ridge^7^ (red lines - Fig 1a, compare similar trajectories between HH27 to HH29 of normal development – black lines). Thus, HH24g mesenchyme (distal 150 micron of graft) exhibits a similar cell cycle profile to HH29 mesenchyme (stage of donor limb), rather than contralateral HH24 mesenchyme (stage of host – Fig 1a - red lines). This could suggest that proliferative outgrowth terminates intrinsically at the end of the patterning phase in the distal mesenchyme independently of e-m signalling (arrows between graft in green and host ridge in purple, Fig. 1b). An alternative possibility is that e-m signalling had irreversibly broken down in donor tissue at the time of grafting and that this led to loss of proliferation. In this scenario, the host apical ectodermal ridge would be extrinsically maintained by host tissue adjacent to the grafts (arrows, Fig. 1c).

**Figure 1.**
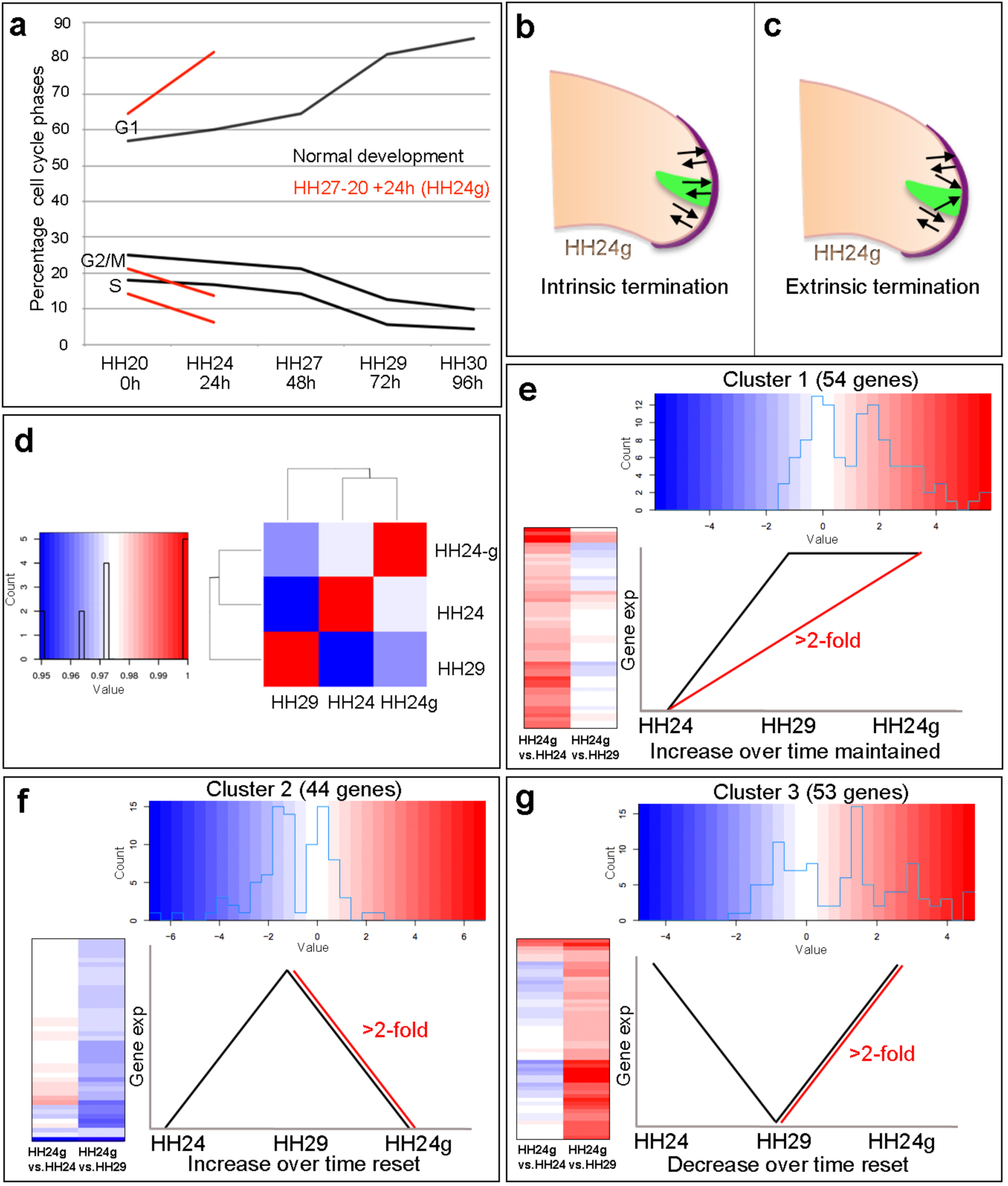
Cell cycle and RNA-seq analyses of chick wing distal tips. a) Decline in cell cycle rate determined by proportions of cells in G1-, S- and G2/M-phases between HH20 and HH30 in the distal chick wing bud (black lines^7^). Red lines show maintenance of cell cycle program in grafts of HH27 distal tips made to HH20 wing buds left for 24 until HH24 (HH24g - tissue would have progressed from HH27 to HH29 in donor) – note trajectories follow same pattern as black lines between HH27 and HH29. **b-c**) Predictions for loss of proliferative growth in HH24g mesenchyme. **b**) Intrinsic termination: e-m signalling (arrows) maintained between graft (green) and apical ridge (purple) and proliferation declines intrinsically in mesenchyme. **c**) Extrinsic termination: e-m signalling lost between graft and apical ridge – note ridge is maintained by signals from host mesenchyme. **d**) Heat-map showing the correlation (Pearson) of the normalised RNA-seq data collapsed to the mean expression per group and the degree of correlation indicated by the colour (red: higher, blue: lower). **e-g**) Clustering of RNA-seq data across pair-wise contrasts with degree of gene expression change indicated by the colour (red: higher, blue: lower). **e**) Cluster 1: genes that increase between HH24 and HH29 (black line) and are maintained in HH24g (red line ->2-fold higher in HH24g than HH24). **f**) Cluster 2: genes that increase between HH24 and HH29 (black line) and reset in HH24g (red line ->2-fold lower in HH24g than HH29). **g**) Cluster 3: genes that decrease between HH24 and HH29 (black line) and reset in HH24g (red line ->2-fold higher in HH24g than HH29).

In this study, we show that wing bud outgrowth at the end of patterning phase terminates intrinsically in the distal mesenchyme in the presence of e-m signalling. Our data provide evidence that an intrinsic switch from BMP antagonism to BMP signalling controls the cell cycle programme in distal mesenchyme during limb bud outgrowth.

## Results

### RNA sequencing reveals reversibility and stability of gene expression

To gain insights into how intrinsic timing is maintained in limb development we used RNA-seq to determine how the transcriptome changes over time in the distal part of the wing bud during normal development and then compared this to the HH24g transcriptome. To achieve this, we performed RNA-seq of pooled blocks of distal cells from HH24, HH24g, HH27 and HH29 chick wing buds (three replicates of 12 blocks taken from the distal mesenchyme excluding the polarizing region– note one HH24 sample failed quality control - see methods). The region of undifferentiated mesenchyme is considered to extend proximally beneath the ridge by 200-300 microns and to remain at a consistent length throughout outgrowth^19^, but for our analyses, we used RNA extracted from the distal 150 microns of control buds and grafts to ensure consistency between all samples. We compared HH24 with HH24g datasets, and HH24g with HH29 datasets (the stage the graft would have developed to if left in situ in the donor wing). This identified 154 genes that are differentially expressed (> 2-fold difference at a *p*-value of 0.0005) between the two comparisons: 55 genes between the HH24g-HH24 pair and 99 genes between the HH24g-HH29 pair (Supplementary datasets 1 and 2). Pearson correlation of the normalised data reveals that overall gene expression levels in HH24g distal cells are closer to HH24 than to HH29 cells, suggesting a good degree of resetting behaviour (Fig. 1d). This is unexpected considering the idea that older mesenchyme cells behave intrinsically in terms of their proliferative and patterning potential when grafted to a younger bud^7^.

To identify genes with similar behaviour across the two pairwise contrasts we performed hierarchical clustering analyses of the 154 genes. This separated the genes into three clusters (See methods – note three genes were excluded that fell into more than one cluster). Cluster 1 (Fig. 1e, Supplementary Fig 1, Supplementary dataset 3) contains 54 genes that increase in expression between HH24 and HH29 during normal development (black line) and are maintained in HH24g distal cells (maintain a >2- fold increase in expression compared to equivalent area in contralateral HH24 buds – red line). Cluster 2 (Fig. 1f, Supplementary Fig 2 and Supplementary dataset 4) contains 44 genes that normally increase in expression between HH24 and HH29 (black line) and are not maintained in HH24g distal cells (>2-fold decrease in expression compared to HH29 buds - red line). Cluster 3 (Fig. 1g, Supplementary Fig 3 and Supplementary dataset 5) contains 53 genes that decrease in expression between HH24 and HH29 (black line) and reset in HH24g grafts (>2-fold increase in expression compared to HH29 buds - red line). Thus genes in cluster 1 reflect stability of expression while genes in cluster 2 and 3 show reversibility of expression. Interestingly, a cluster containing genes whose decrease in expression over time is maintained in HH24g distal cells was not found indicating that the young environment potently reactivates genes that decrease in expression. These data suggest that many genes preferentially expressed during early stages of wing development can be reset to some degree (Cluster 3). However, later expressed genes show differential resetting behaviour in an earlier environment (Clusters 1 and 2).

**Figure 2.**
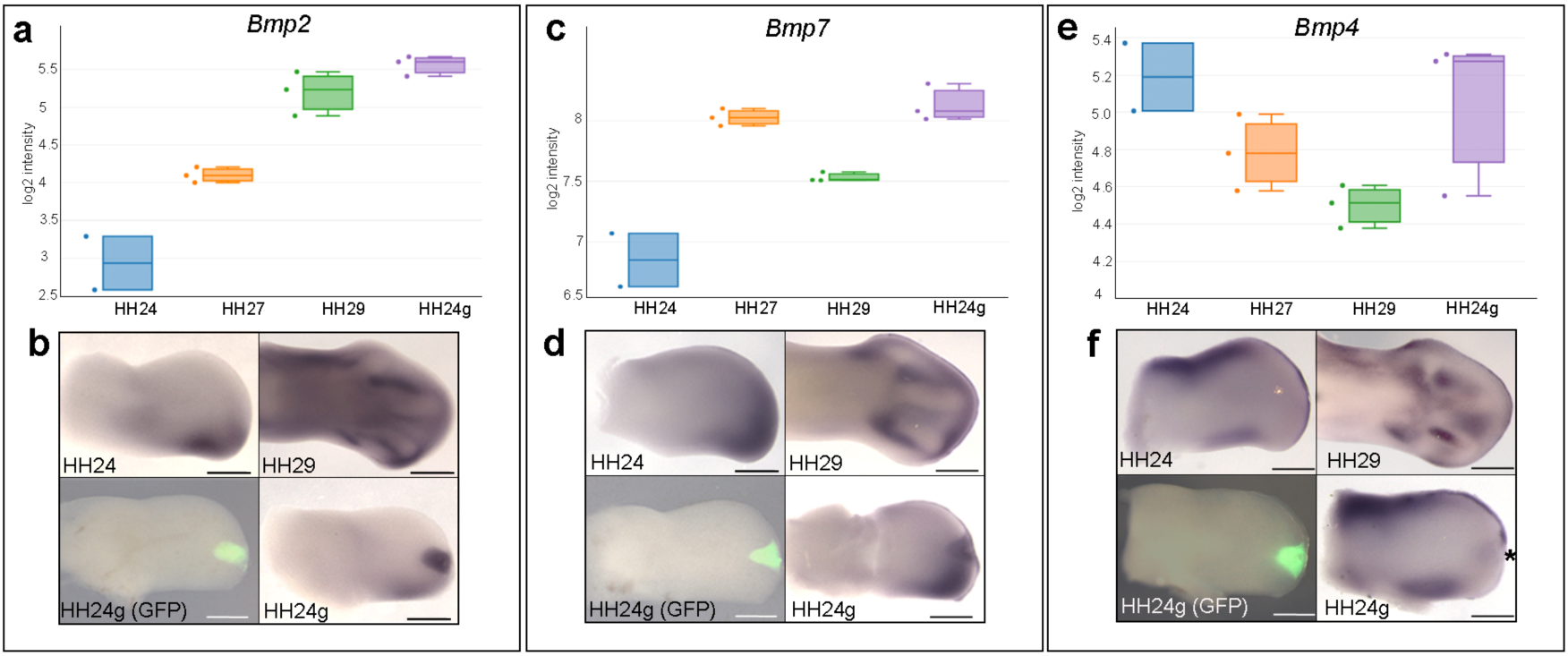
Maintenance of *Bmp2/4/7* expression in HH24g mesenchyme. a) Histogram showing expression levels of *Bmp2* as normalised log_2_ values of RNA sequencing read-count intensities. **b**) In situ hybridization showing *Bmp2* expression - note intense expression in HH24g mesenchyme in area of graft (*n*=3/3, HH24 bud is the contralateral). **c**) Expression levels of *Bmp7* determined by RNA-seq. **d**) In situ hybridization showing *Bmp7* expression - note intense region of expression in HH24g mesenchyme (*n*=3/3). **e**) Expression levels of *Bmp4* determined by RNA-seq. **f**) In situ hybridizations showing *Bmp4* expression - note enhanced expression in HH24g mesenchyme in area of graft (*n*=2/3). Scale bars: HH24 buds - 500µm; HH29 buds - 200µm.

**Figure 3.**
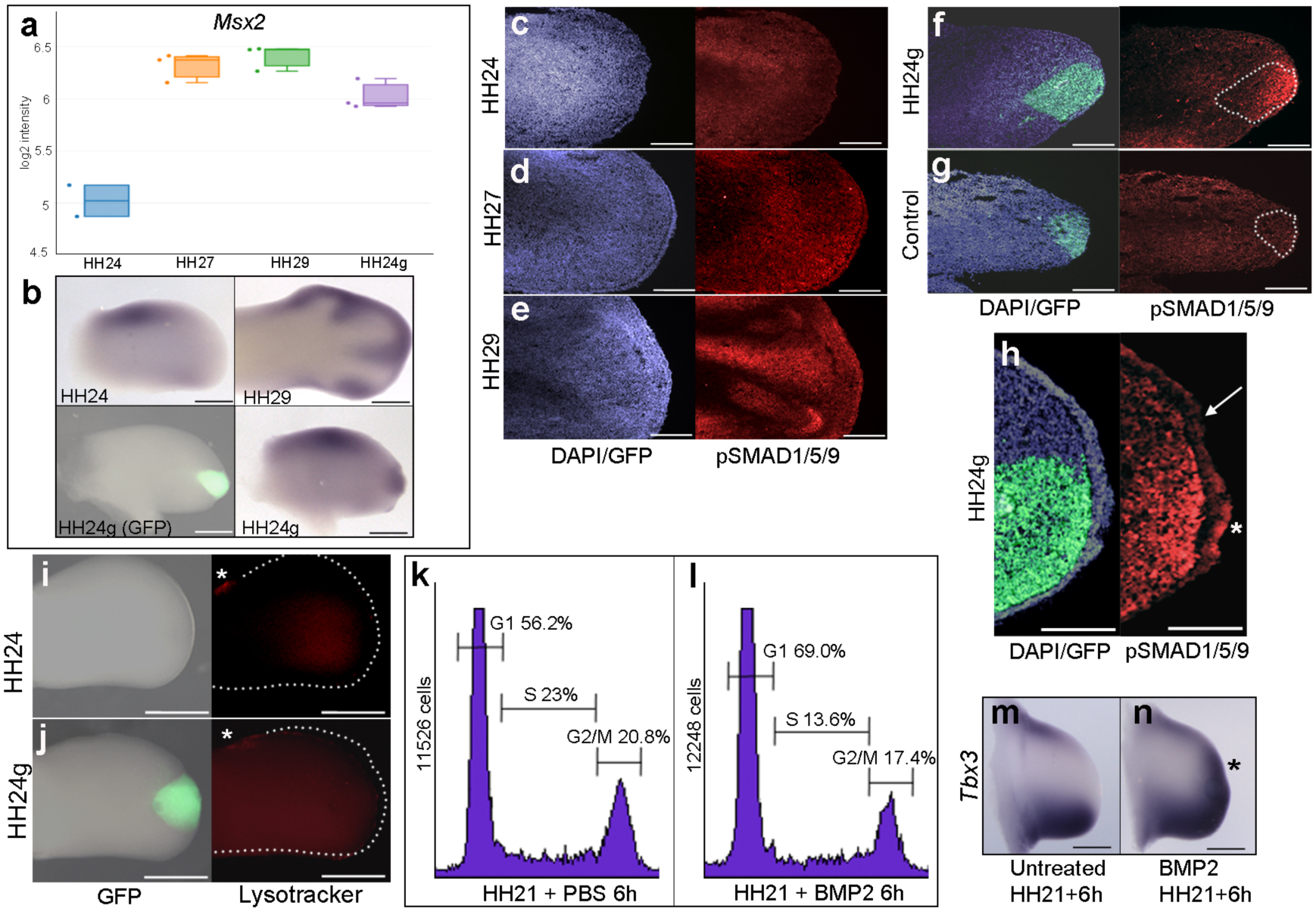
Maintenance of BMP signalling in HH24g mesenchyme. a) Expression levels of *Msx2* determined by RNA-seq. **b**) In situ hybridization showing *Msx2* expression - note intense expression in HH24g mesenchyme in grafted area (*n*=3/4). **c-h**) Immunostaining of pSMAD1/5/9: signal increases in the distal part of chick wing buds over time (**c-e**) and is enhanced in HH24g mesenchyme (**f**) compared to HH24 buds (*n*=5/5 - **c**) but not in HH20-HH20 grafts (*n*=2/2 - **g**); signal is enhanced in apical ectodermal ridge above grafts in HH24g mesenchyme (asterisk-**h**) but not adjacent regions of ridge (*n*=2/2 - arrow-**h**). **i-j**) Lysotracker staining of apoptotic cells reveals no difference in HH24 (**i**) and HH24g (**j**) mesenchyme (*n*=5/5) – asterisks indicate anterior necrotic zone. **k-l**) Flow cytometry of wing bud distal mesenchyme treated for 6 h with PBS-soaked bead at HH21 showing cell cycle parameters (**k**) and significant increase in proportion of cells in G1-phase and decrease in proportions in S- and G2/M-phases (Pearson’s χ^2^ test - *p*<0.0001) after 6h BMP2-soaked bead treatment (**l**). **m-n**) Expression of *Tbx3* in chick wing (**m**) and enhanced expression in proximity of a BMP2-soaked bead implanted at HH21 after 6h (*n*=4/4 – asterisk in **n**). Scale bars: HH24 buds - 500µm, HH29 buds - 200µm in b; 150µm in c-g; 75µm in h; 300µm in i-j; 500µm in m-n.

### Bmp2/4/7 expression is maintained in HH24g mesenchyme

To identify signalling pathways associated with the maintenance of intrinsic cell cycle timing in HH24g mesenchyme, we focused on cluster 1 that contains genes expressed at late stages of development and are maintained in distal mesenchyme transplanted to an earlier environment (Fig. 1e, Supplementary Fig 1, Supplementary dataset 3). This cluster notably includes genes encoding members of the Bone Morphogenetic Protein (BMP) family - *Bmp2* and *Bmp7*. Genes encoding other known signalling molecules are not present in cluster 1. BMP signalling is implicated in terminating outgrowth during the patterning phase by inhibiting FGF signalling by the apical ectodermal ridge^16^. Read-counts of the RNA-seq data show that the expression of *Bmp2* increases by 5.3-fold between HH24 and HH29 – similar to the difference in expression in HH24g tissue compared to HH24 tissue (6.2-fold - log_2_ scale is shown), thus showing that late expression levels of *Bmp2* are maintained in an earlier environment (Fig 2a). Similarly, a 1.5-fold increase occurs with *Bmp7* between HH24 and HH29 and the difference between expression levels in HH24g tissue compared to HH24 tissue is 2.5-fold (Fig. 2c). Analyses by in situ hybridization are consistent with these observations (Figs 2b and 2d). We also analysed the expression of *Bmp4*, which is not present in a cluster, but has a prominent role in limb development^20 21^. The RNA-seq data indicates that *Bmp4* expression levels decrease by 2-fold between HH24 to HH29 and that expression levels are approximately equivalent between HH24 and HH24g tissue (Fig.2e, note stronger expression in ridge at HH24 compared to *Bmp2/7*). Despite this, slightly enhanced expression could be detected by in situ hybridization in the region of the grafted tissue in comparison to the equivalent position in contralateral buds (asterisk - Figure 2f).

### BMP signalling is maintained in HH24g mesenchyme

To determine if *Bmp* expression in HH24g mesenchyme is associated with an increase in transcription of BMP target genes, we analysed the expression of *Msx2^22^*, which is not present in a cluster. RNA-seq data for *Msx2* reveals a similar profile to both *Bmp2* and *Bmp7*: a 2.8-fold increase between HH24 and HH29 and a 2-fold difference in HH24g tissue compared to HH24 tissue (Fig. 3a). RNA in situ hybridization of *Msx2* confirms this result (Fig 3b). Consistent with this finding, immunofluorescence using an antibody that recognizes phosphorylated SMAD1/5/9 – the signal transducer for BMPs – reveals increased levels in sections of later wing buds (Figs 3c-e.). However, whereas weak expression is detected in HH24 wing buds (Fig. 3c), strong expression is observed in the distal part of grafts (the area processed for RNA-seq) in contralateral HH24g buds (Fig. 3f). Control grafts of HH20 distal mesenchyme made to host HH20 wing buds show no obvious differences in pSMAD1/5/9 signal intensity after 24 h compared to stage-matched HH24 buds (Fig. 3g–compare to Fig. 3c). Interestingly, enhanced pSMAD1/5/9 signal is detected in the host ridge above HH24g mesenchyme (asterisk Fig. 3h), in contrast to adjacent regions (arrow – Fig 3h). This result suggests that the level of BMP signalling is maintained in mesenchyme transplanted from older buds to a younger environment, but is also enhanced in the apical ectodermal ridge in response to the older mesenchyme.

BMP signalling is associated with regions of apoptosis in the developing limb, including anterior and posterior necrotic zones, as well as the interdigital mesenchyme^23^. To determine if apoptosis could reduce proliferation in HH24g mesenchyme, we performed lysotracker staining of apoptotic cells. This reveals no appreciable difference in HH24 control (Fig. 3i) and HH24g mesenchyme (Fig. 3j) in whole-mount preparations. Note, the anterior necrotic zone can be observed in both examples (asterisks in Fig. 3i and 3j).

To test directly the effects of BMP signalling on cell-cycle dynamics in early wing buds, we implanted beads soaked in recombinant BMP2 protein into the distal mesenchyme of HH21 buds. We used flow cytometry to determine the proportion of cells in the different phases of the cell cycle in distal tissue (150 microns from the distal tip) of HH21 wing buds that were treated either with BMP2-soaked beads or PBS-soaked beads for 6 h. In wing buds treated with BMP2, the proportion of cells increases in G1-phase by 13% (56-69%), but decreases in S-phase by 9% (23-14%), and in G2/M-phase by 4% (21-17% - Fig. 3k and 3l). Enhanced expression of *Tbx3* confirms BMP2 activity in cells around the beads after 6 h (Fig. 3m and asterisk in Fig. 3n). This finding shows that enhanced BMP signalling reduces proliferation of distal mesenchyme cells in the early chick wing bud. Taken together these results suggest that BMP signalling in HH24g mesenchyme is involved in maintenance of an intrinsic cell cycle programme.

### e-m signalling is reset in HH24g mesenchyme

The maintenance of BMP signalling in HH24g mesenchyme could reduce proliferation via inhibition of FGFs produced by the apical ectodermal ridge. However, we have previously shown that *Fgf8* expression is maintained in the ridge above grafts of distal mesenchyme from much older wing buds - including in tissue that would have developed to stage HH36 if left in situ^7^ (or day 10 of incubation – the ridge normally regresses at HH29/30 in the chick wing at day 6-6.5 of incubation). Another possibility is that HH24g cells have progressed beyond a point at which they irreversibly fail to respond to FGFs derived from the apical ectodermal ridge. To address this, we looked at expression of a transcriptional target of FGF signalling, *Dusp6 ^24^* (also known as *Mkp3* and *Pyst1*). Although not in one of the clusters, RNA-seq read-counts show that *Dusp6* expression reduces by 2.1-fold between HH24 and HH29, but that no significant difference in expression is observed between HH24g tissue and the equivalent region of contralateral control buds, indicating that the level of FGF signalling is the same in the grafted cells as in the host wing bud (Figs 4a and b). In addition, cluster 2 contains *Fgfr2*, which encodes an FGF receptor that is expressed in the distal mesenchyme and apical ectodermal ridge^25^– note cluster 2 contains few other well-known genes involved in limb development (Supplementary Fig 2, Supplementary dataset 4). RNA-seq read-counts reveal that expression of *Fgfr2* increases 2.8-fold between HH24 and HH29, but is approximately the same in HH24 and HH24g tissue (a 1.2-fold increase in HH24g – Figs 4c and d – note that *Fgfr1* expression, which is also expressed in distal mesenchyme and can respond to ridge signals ^25^, is approximately the same in HH24, HH24g and HH29 buds **–** not shown).

**Figure 4.**
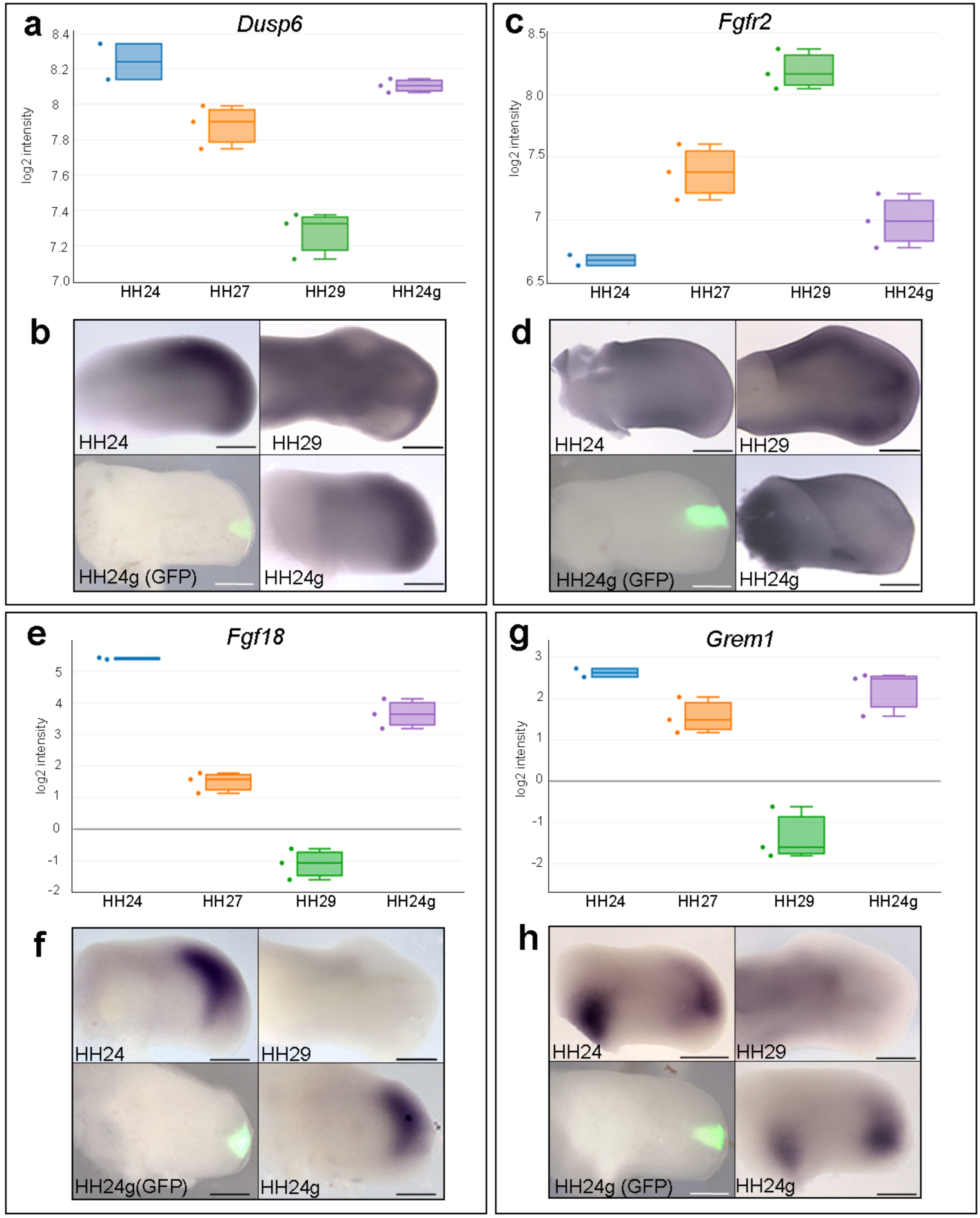
Resetting of e-m signalling in HH24g mesenchyme. a-h) Resetting of gene expression in HH24g mesenchyme. Expression levels of *Dusp6* (**a**)*, Fgfr2* (**c**)*, Fgf18* (**e**) and *Grem1* (**g**) determined by RNA-seq. **b**) In situ hybridization showing equivalent levels of *Dusp6* (*n*=3/3 **b**)*, Fgfr2* (*n*=4/4, **d**)*, Fgf18* (*n*=6/7, **f**) and *Grem1* (*n*=7/9, **h**). Scale bars: HH24 buds - 500µm; HH29 buds - 200µm.

Cluster 3 contains genes encoding other members of the FGF family, including *Fgf13* and *Fgf18*, which are expressed in distal mesenchyme at early stages^26^ ^27^ and that can be reset in a younger environment (Supplementary Fig 3, Supplementary data 5). RNA-seq read-counts show that *Fgf18* decreases by 90-fold between HH24 and HH29, by which point it is undetectable by in situ hybridization (Figs 4e and f), but expression is only 3.4-fold higher in HH24 tissue compared to HH24g tissue. In addition, cluster 3 also contains *Grem1*, which encodes the BMP antagonist that maintains the apical ectodermal ridge (Figs 4l and m, Supplementary Fig 3, Supplementary data 5). Between HH24 and HH29, *Grem1* expression levels are reduced by 17.2-fold until they are almost undetectable by in situ hybridization (Figs 4g and h). However, *Grem1* expression is approximately the same in HH24 and HH24g mesenchyme (Figs 4g and h). Taken together, these results show that proliferation of HH24g mesenchyme cells declines in the presence of e-m signalling.

## Discussion

By grafting late mesenchyme cells into an early epithelial environment, we have determined the function of e-m signalling in the decline and cessation of chick wing bud outgrowth at the end of the patterning phase. Our data reveals that the duration of mesenchyme cell proliferation is intrinsically determined, but requires permissive signals from the apical ectodermal ridge: a distinction not recognized in previous approaches. For instance, when apical ectodermal ridge signalling is disrupted by embryological approaches in the chick wing^12^ ^13^, or by genetic approaches in the mouse limb^28^, mesenchyme expansion is naturally inhibited, thus masking the intrinsic control of proliferation in this tissue.

A schematic detailing the relationship between e-m signalling and mesenchyme proliferation during normal wing outgrowth is shown in Figure 5. At early stages of wing development (HH20), the Grem1-mediated inhibition of BMP signalling fulfils two main roles: to maintain the apical ectodermal ridge as classically-described; and to permit a rapid rate of proliferation in distal mesenchyme cells as we have shown here. Around HH20, the mechanism of positional specification switches in mesenchyme progenitors from extrinsic signalling (humerus)^4 10^ to an intrinsic timer (distal to elbow)^6^. This intrinsic mechanism involves a progressive reduction in the proliferation rate as the elements are laid down in a proximal to distal sequence (Fig. 5). The deceleration in proliferation rate is caused by an increase in BMP signalling between HH24 and HH29 that overcomes Grem1-mediated antagonism in distal mesenchyme cells. Once the intrinsic cell cycle timer has run its course and distal mesenchyme expansion ceases, high BMP signalling causes the apical ectodermal ridge to regress at HH29/30 (Fig 5). Therefore, BMP signalling forms an autoregulatory circuit in which it controls a cell cycle timer in the mesenchyme, while extrinsically maintaining the apical ectodermal ridge to permissively allow the timer to run.

**Figure 5.**
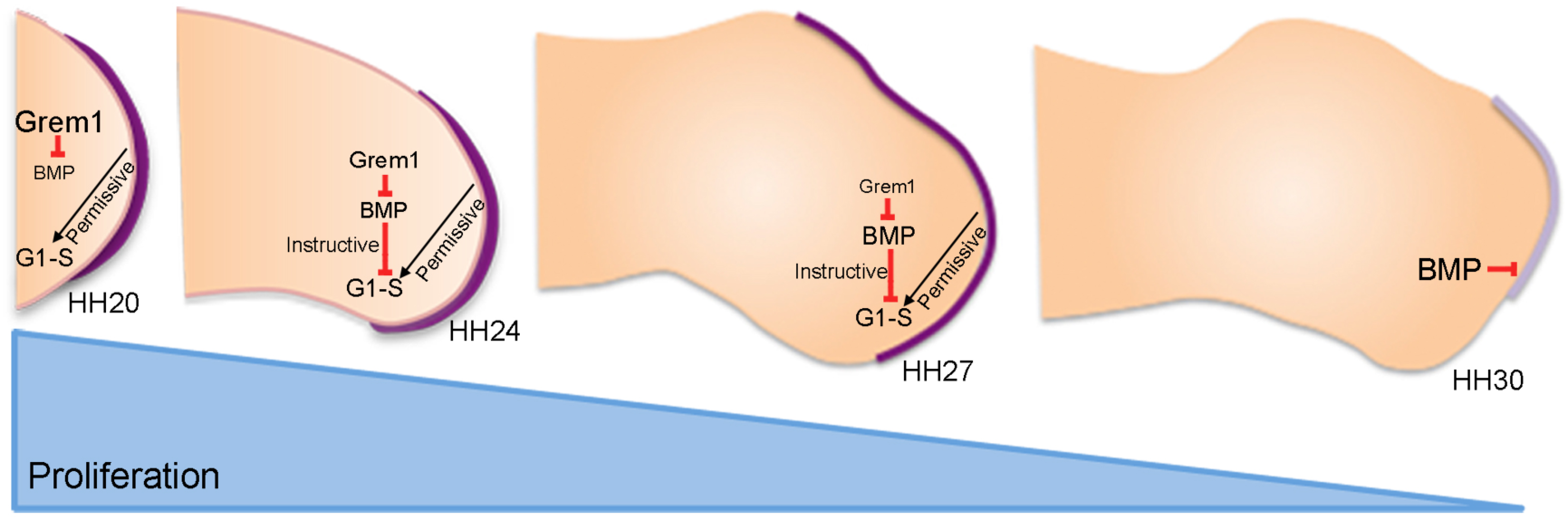
An intrinsic cell cycle timer terminates limb bud outgrowth. HH20-HH24: antagonism of BMP signalling by Grem1 permits rapid mesenchyme proliferation and maintenance of the apical ectodermal ridge (purple) that permissively supports outgrowth. HH24-HH27: intrinsic rise in BMP signalling counters BMP antagonism and instructively decelerates mesenchyme proliferation rate. HH27-HH30: mesenchyme proliferation ceases and then BMP signalling inhibits FGF signalling in apical ectodermal ridge that then regresses.

It was unexpected that all the genes identified in the RNA-seq analysis that are specifically expressed at early stages could be reset to some degree by the signalling environment. This is consistent with the observations that early proximo-distal patterning is controlled by extrinsic signals – possibly retinoic acid signalling from the flank of the embryo^4 10^. Notably, this cluster contains several genes encoding products known to be involved in maintaining a stem cell-like state including *Plzf* (*Zbtb16*)^29^ *Sall4*^30^ (also *Sall1* and *3*), *Gbx2*^31^ and *Lin28b*^32^. This raises the possibility that a gene regulatory network operates during early limb development to maintain the multipotency of mesenchyme cells and warrants further investigation.

Opposing gradients of BMP ligands and their antagonists specify positional identities and regulate growth in a concentration-dependent manner in many developmental contexts^33^. Thus, the intrinsic switch from BMP antagonism to BMP signalling that we describe here provides a mechanism by which cells can be instructed by defined concentrations of BMP signalling in the absence of spatial gradients. It will be important to identify other patterning systems that use a similar mechanism.

## Methods

### Chick husbandry and tissue grafting

Wild type and GFP-expressing Bovans Brown chicken eggs were incubated and staged according to Hamburger Hamilton^34^. For tissue grafting the polarizing region was discarded and a stripe of 150 µm of distal sub-AER mesenchyme about 500 µm long was dissected from HH27 wing buds. The ectoderm was removed after incubation in 0.25% trypsin at room temperature for 2 mins and the mesenchyme was then cut into three cuboidal pieces. These were then placed in slits made using a fine sharpened tungsten needle between the apical ectodermal ridge and underlying mesenchyme in the mid-distal region of HH20 wing buds.

### RNA sequencing analyses and clustering

Grafts were made as described above and after 24 h the 150 µm distal part of the graft was carefully dissected under a UV scope. Equivalent tissue was dissected from the contralateral HH24 wing bud and also 150 µm of tissue was dissected from three positions at the distal tips of HH24, HH27 and HH29 wing buds. Three replicate experiments were performed from each condition and the tissue was pooled before the RNA was extracted using Trizol reagent (Gibco), and sequenced on a HiSeq 2000(Paired end readings of 50bp - Instrument: ST300). Sequencing data was mapped using Hisat v2.0.3 (bam file generation). Salmon output was used to quantify the data (number of reads)^35^. The raw data has been deposited in array express (https://www.ebi.ac.uk/arrayexpress/experiments/E-MTAB-6437/). 12 samples (three for each of the four conditions HH24, HH24g, HH27 and HH29) were checked for quality issues. This was done by manually inspecting the density plot, boxplots, PCA plots, correlation heatmap and distance plot, as well as using several automatic outlier tests, namely distance, Kolmogorov-Smirnov, correlation and Hoeffding’s D. Based on this, one of the HH24 samples was excluded from the analysis.

The count data for the samples were normalised using trimmed mean of *m*- value normalisation and transformed with Voom, resulting in log_2_-counts per million with associated precision weights^36^ ^37^. A heat-map was made showing the correlation (Pearson) of the normalised data collapsed to the mean expression per group. A statistical analysis using an adjusted *p*-value < 0.0005 and a fold-change >2 identified 151 genes as differentially expressed in the two contrasts evaluated. Gene clusters were identified from the set of 151 unique genes that were differentially expressed in at least one of the statistical comparisons. The evaluation considered between two and 35 clusters using hierarchical, *k*-means, and PAM clustering methods based on the internal, stability and biological metrics provided from the clValid R package. The majority of internal validation and stability metrics indicated that either the lowest possible number or conversely the highest number evaluated were preferable. The metrics giving more nuanced information in the intermediate range were the Silhouette measure, and the Biological Homogeneity Index (BHI). Based on manual inspection it was decided to use hierarchical clustering with *k*=3 gene clusters, which showed favourable properties for both these measures.

### Apoptosis analyses

Chick wing buds were dissected in PBS and transferred to Lysotracker (Life Technologies, L-7528) PBS solution (1:1000) in the dark. Wing buds were incubated for 1 h at 37°C, washed in PBS, and fixed overnight in 4% PFA at 4°C. Wing buds were then washed in PBS and progressively dehydrated in a methanol series.

### Whole mount in situ hybridisation

Embryos were fixed in 4% PFA overnight at 4°C, dehydrated in methanol overnight at −20°C, rehydrated through a methanol/PBS series, washed in PBS, then treated with proteinase K for 20 mins (10µg/ml-1), washed in PBS, fixed for 30 mins in 4% PFA at room temperature and then prehybridised at 69°C for 2 h (50% formamide/50% 2x SSC). 1µg of antisense DIG-labelled mRNA probes were added in 1 ml of hybridisation buffer (50% formamide/50% 2x SSC) at 69°C overnight. Embryos were washed twice in hybridisation buffer, twice in 50:50 hybridisation buffer and MAB buffer, and then twice in MAB buffer, before being transferred to blocking buffer (2% blocking reagent 20% lamb serum in MAB buffer) for 2 h at room temperature. Embryos were transferred to blocking buffer containing anti-digoxigenin antibody (1:2000) at 4°C overnight, then washed in MAB buffer overnight before being transferred to NTM buffer containing NBT/BCIP and mRNA distribution visualised using a LeicaMZ16F microscope.

### Immunohistochemistry

Embryos were fixed in 4% PFA for 2 h on ice, washed in PBS and transferred to 30% sucrose overnight at 4°C. Embryos were frozen in OCT mounting medium, fixed to a chuck and cryosectioned immediately. Sections were dried for 2 h and washed in PBS containing 0.1% triton for 10 mins at room temperature. Sections were blocked in 0.1% Triton, 1-2% hings/serum for 1-1.5 h at room temperature. Primary antibody solution was added (1:500 rabbit anti-PSMAD1/5/9 - Cell Signalling Technology – D5B10) in PBS /0.1% triton/1-2% hings, before coverslips were added and slides placed in a humidified chamber at 4°C for 72 h. Slides were then washed for 3x 5mins in PBS at room temperature. Secondary antibody (1:500 anti-rabbit Alexa Fluor 594 - Cell Signalling Technology – 8889S) was added in PBS 0.1% triton/1-2% hings for 1 h at room temperature. Slides were washed in PBS for 3x 5 mins and mounted in Vectashield/DAPI medium. Slides were left at 4°C and imaged the following day.

### Flow cytometry

Distal 150 µm blocks of mesenchyme tissue were dissected in ice cold PBS under a LeicaMZ16F microscope using a fine surgical knife pooled from replicate experiments (12), and digested into single cell suspensions with trypsin (0.5%, Gibco) for 30 mins at room temperature. Epithelium was removed and cells were briefly washed twice in PBS, fixed in 70% ethanol overnight, washed in PBS and re-suspended in PBS containing 0.1% Triton X-100, 50µg/ml^-1^ of propidium iodide and 50µg/ml^-1^ of RNase A (Sigma). Dissociated cells were left at room temperature for 20 mins, cell aggregates were removed by filtration and single cells analysed for DNA content with a FACSCalibur flow cytometer and FlowJo software (Tree star Inc). Based on ploidy values cells were assigned in G1, S, or G2/M phases and this was expressed as a percentage of the total cell number (>11,000 cells in each case). Statistical significance of numbers of cells between pools of dissected wing bud tissue (12 in each pool) was determined by Pearson’s *χ*^2^ tests to obtain two-tailed *P*-values (significantly different being a *P*-value of less than 0.05 - see^38^ - statistical comparisons of cell cycle parameters between the left and right wing buds embryos showed that differences are less than 1%).

### Bead implantation

Affigel blue beads (Biorad) were soaked in PBS or recombinant human BMP2 protein (150µg/ml^-1^ R&D) dissolved in PBS/4mM HCl. Beads were soaked for 2 h and implanted into the distal mesenchyme using a sharp needle.

## Acknowledgements

We thank the Wellcome Trust for funding, Cheryll Tickle for critical reading, Adrian Sherman/Helen Sang for GFP-expressing chicken embryos, the University of Sheffield flow cytometry core facility and Max Bylesjo at FIOS Genomics for bioinformatics.

## Competing interests

The authors declare no competing interests

## Author contributions

JP performed/planned all experiments, processed samples for RNA-seq and wrote early drafts, KC performed grafts for RNA-seq, CR contributed to RNA in situ analyses, P. S-L performed pilot experiments, MR performed pilot experiments and edited the paper, MT planned the research and wrote the paper.

## Funding

The authors were supported by the Wellcome Trust (202756/Z/16/Z) and by the Spanish Ministerio de Economía, Industria y Ccompetitividad (BFU2017-88265-P).

